# Himalayan whole-genome sequences provide insight into population formation and adaptation

**DOI:** 10.1101/2024.11.26.625458

**Authors:** Elena Arciero, Mohamed A. Almarri, Massimo Mezzavilla, Yali Xue, Pille Hallast, Cidra Hammoud, Yuan Chen, Laurits Skov, Thirsa Kraaijenbrink, Qasim Ayub, Huanming Yang, George van Driem, Mark A. Jobling, Peter de Knijff, Chris Tyler-Smith, Asan, Marc Haber

## Abstract

High-altitude environments pose substantial challenges for human survival and reproduction, attracting considerable attention to the demographic and adaptive histories of high-altitude populations. Previous work focused mainly on Tibetans, establishing their genetic relatedness to East Asians and their genetic adaptation to high altitude, especially at *EPAS1*. Here, we present 87 new whole-genome sequences from 16 Himalayan populations and the insight they provide into the genomic history of the region. We show that population structure in the Himalayas began to emerge as early as 10,000 years ago, predating archaeological evidence of permanent habitation above 2,500 meters by approximately 6,000 years. The high prevalence of the introgressed adaptive *EPAS1* haplotype in all high-altitude populations today supports a shared genetic origin and its importance for survival in this region. We also identify additional selection signals in genes associated with hypoxia, physical activity, immunity and metabolism which could have facilitated adaptation to the harsh environment. Over time, increasing genetic structure led to the diverse and strongly differentiated ethnic groups observed today, most of which maintained small population sizes throughout their history or experienced severe bottlenecks. Between 6,000 and 3,000 years ago, a few uniparental lineages became predominant, likely coinciding with the advent of agriculture, although significant population growth was not observed in the Himalayas except in the Tibetans. In more recent times, we detect bidirectional gene flow between high-altitude and lowland groups, occurring on both sides of the Himalayan range. The timing of this admixture aligns with the rise and expansion of historical regional powers, particularly during the Tibetan Empire and the northern Indian Gupta Empire. In the past few centuries, migrations to the Himalayas seem to have occurred alongside conflicts and population displacements in nearby regions and show some sex bias.

## Introduction

Even for brief visits, high altitudes are challenging for most humans because of factors including the low oxygen concentration and cold temperatures. Living at high altitude throughout the year introduces additional requirements for sources of food and other essentials that can be produced there^1^. High-altitude living has therefore been the focus of intensive investigation, particularly in the Himalayas, where many present-day populations live permanently above 3,000 meters (m). Information relevant to understanding the history of these populations and the adaptations they have undergone comes from diverse fields, including fossils, archaeology, linguistics and genetics. The earliest known high-altitude Himalayan hominin fossil is a 160,000-year-old mandible from Tibet identified as Denisovan by protein analysis^2^. Evidence for modern human presence dates back to 30,000-40,000 years ago (ya), when the Nya Deḥu site at 4,600 m in Tibet shows archaeological signs of occupation in the form of stone tool production^3^. Nevertheless, a strong case has been made that villages for year-round settlement at altitudes up to 2,500 m were only established after 5,200 ya when millet could be cultivated, and that permanent habitation up to 3,500 m only occurred after 3,600 ya when barley and sheep from the west became available^4^. Current high-altitude Himalayan populations mostly speak Bodic languages, and a lexicostatistical study has generated a glottochronological estimate for Bodic, including Tamangic and the linguistic subgroups formerly covered by the label ‘East Bodish’ at between 4,000 and 3,000 ya^5^.

Genetic analyses of present-day populations, mostly focusing on Tibetans, have contrasted the diverse mtDNA lineages with the limited Y-chromosomal lineages^6,7^, emphasised the range of reported coalescence times within the Tibetan population from 7,400-29,900 ya, and suggested that Tibet could have been initially populated since 30,000 ya with additional gene flow occurring 7,000-10,000 ya^7^. Whole-genome sequencing-based analyses have demonstrated the overall genetic relatedness of Tibetans to East Asian populations, splitting from Han ∼15,000-9,000 ya^8^, or 44,000-58,000 ya with extensive gene flow until 9,000 ya^9^. These contrast with an estimate for the Tibetan-Han split from exome sequences of 2,750 ya^10^. Ancient DNA studies in the region, while relatively limited, reveal a complex Tibetan ancestry that derives mostly from 4,000 to 9,000 year-old populations in northern East Asia^11^. Additionally, samples from the Annapurna Conservation Area in Nepal, dating between 1,250 to 3,150 years old and found at elevations of 2,800 to 4,000 m, exhibited the closest genetic affinity to present-day Tibetans among the examined populations, implying population continuity at moderate to high altitudes^12^.

Studies of genetic adaptation to high altitude initially identified selection at *EPAS1*^10^ in a region that had introgressed from Denisovans^13^, a finding that has been replicated in multiple studies, which have often pointed to additional selected loci^8,9,14^. The selection in Tibetans at Denisovan-introgressed segments stands as one of the strongest examples of beneficial archaic admixture in modern humans. Nevertheless, there has been limited exploration in non-Tibetan populations regarding *EPAS1* or similar signals in other regions of the genome.

Here, we present 87 new high-coverage genome sequences derived from 16 distinct Himalayan populations. We use the new data to study the demographic processes underlying human presence in the Himalayas more widely, addressing previously reported discrepancies and providing details regarding population emergence, expansion timelines, genetic structure, relationships to neighbouring populations and local adaptations.

## Results

### Dataset

We generated high-coverage whole-genome sequences from 87 samples of 16 populations from Bhutan, Nepal, India and the Tibetan plateau (Table S1, Figures 1A-C).

**Figure 1.**
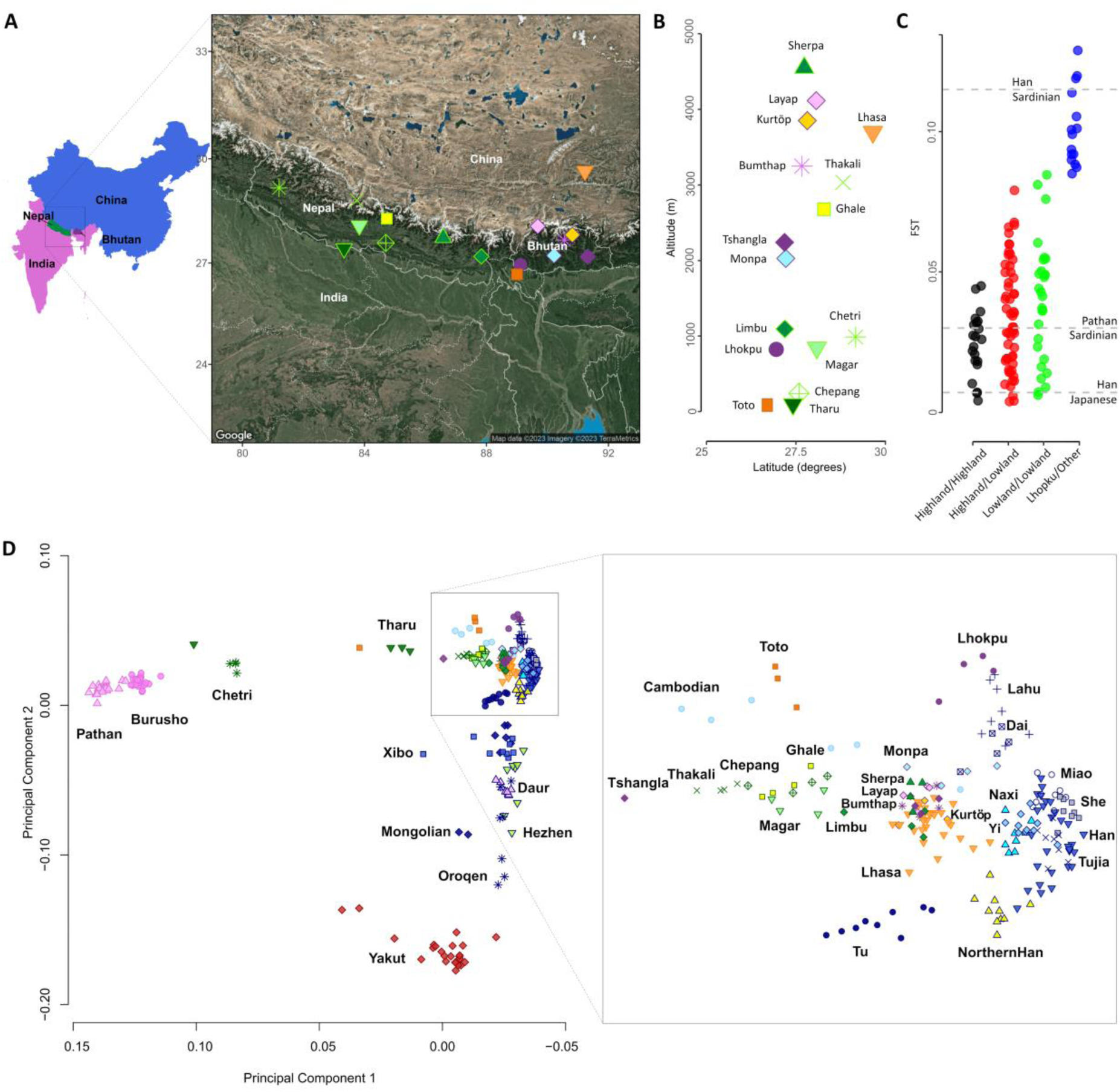
Populations sequenced in this study and their genetic relationships to neighbouring populations. **A)** The newly generated dataset includes 16 populations from China, Bhutan, Nepal and India. Map shows the geographic locations of the new samples. **B)** The plot displays the latitude and altitude of the new populations. **C)** Pairwise *F*ST comparisons between populations. We refer to populations living over 2,500 as ‘Highland’. **D)** Principal component analysis showing the Himalayan populations in a regional context with Central/Southern and East Asian populations. The inset provides a closer view of the main Himalayan cluster.

We performed joint calling of these samples together with the Human Genome Diversity Project (HGDP) dataset^15^ for autosomal, Y-chromosomal and mtDNA sequences. We identified ∼19 million variants in the Himalayan populations, comprising 16 million SNPs and 3 million Indels. Notably, this dataset includes 5 million variants absent from the HGDP dataset, with 2.7 million variants having a Minor Allele Frequency (MAF) > 1%. This highlights the significance of incorporating diverse populations to capture a broader spectrum of human genetic diversity.

### Himalayan genetic structure and relationship to regional populations

Principal Components Analysis (PCA) shows that most of the Himalayan populations cluster together and are close to East Asian Chinese populations (Figure 1D). We detect a genetic structure that distinguishes the Bhutanese populations, as well as the Sherpa and Tibet populations, from the other Nepalese populations - Thakali, Ghale, Chepang and Magar. The latter groups appear relatively drawn toward Central/South (CS) Asian populations. Sherpa and Tibetans overlap and cluster with the Bhutanese high-altitude populations Layap and Kurtöp while the Nepalese high-altitude populations Thakali and Ghale cluster together. The Chetri and Tharu populations from Nepal appear significantly drawn towards CS Asian populations, and the Indian Toto population clusters close to Cambodians. The closest lowland Chinese populations to the Himalayans are Tu, Naxi, Yi and Northern Han.

To further examine the extent of population differentiation, we calculated *F*ST values across 1.3 million SNPs identified as polymorphic in archaic genomes, allowing for unbiased estimates^15^. We find notably high *F*ST values in most pairwise comparisons relative to the geographic distances between populations (Figure 1C; Supplementary Figure 1), with several pairwise estimates exceeding those typically observed between intercontinental groups within Eurasia. The Lhokpu population stands out as an extreme outlier, showing strong differentiation from all other populations, with most *F*ST values exceeding 0.1, and the Tshangla appearing the least differentiated to them (*F*ST 0.085).

As runs of homozygosity (ROHs) can provide insights into demographic history^16^, we identified ROHs greater than 1 Mb in our dataset (Figure 2A, Supplementary Figure 2A). Our findings reveal that the Lhokpu, and to a lesser extent the Monpa, Toto and Chepang, exhibit remarkably high total sums of ROHs per individual (sROHs, 200–500 Mb). Compared to the HGDP dataset, which includes over 50 diverse worldwide populations, these Himalayan populations stand out as having some of the highest sROHs observed globally (Supplementary Figure 2B). The Lhokpu, in particular, show a pattern similar to that of Native South American populations, such as the Surui and Karitiana, which are known to have undergone significant bottlenecks. While these elevated sROHs might initially suggest recent inbreeding, this does not appear to be the primary cause. These Himalayan populations also possess high counts of individual ROH segments (nROHs), which distinguishes them from South Asian and Middle Eastern populations where consanguinity is common. In those consanguineous groups, the pattern typically includes fewer but larger ROHs, indicative of recent inbreeding^17^ (Supplementary Figure 2B). In contrast, these Himalayan groups show a distinct pattern with higher nROHs and lower average ROH sizes, suggesting that these populations experienced ancient bottlenecks, possibly coupled with recent consanguinity. Other populations appear to have comparably very low numbers of ROHs, such as the Lhasa and the Chetri (Supplementary Figure 2A), indicating historically large population sizes, and/or possible admixture.

**Figure 2.**
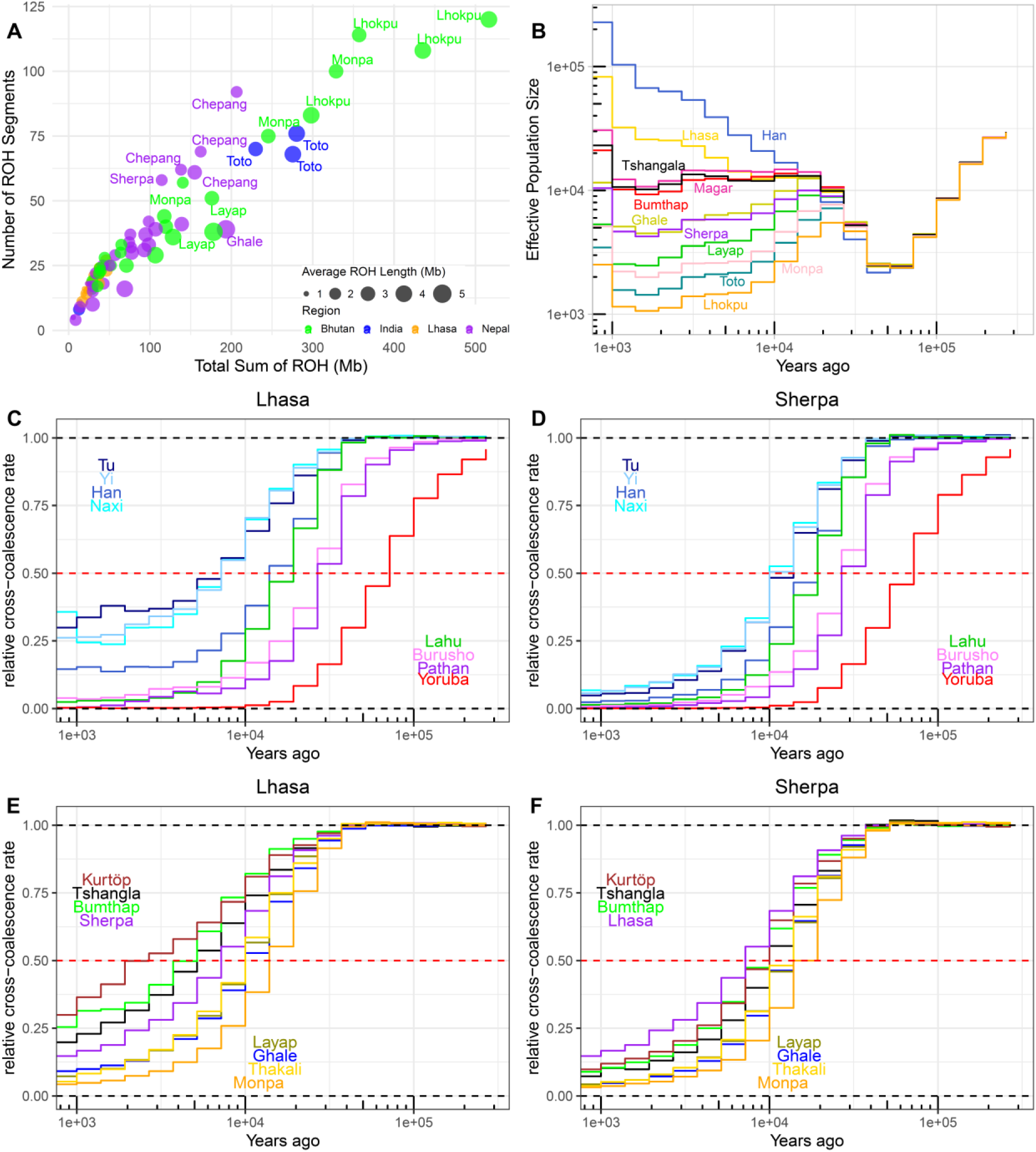
Himalayan population dynamics and divergence times. **A)** Distribution of Runs of Homozygosity in Himalayan populations. **B)** Inferred effective population size (Ne) over time. **C)** Inferred cross-coalescent rate over time between the Lhasa and other non-Himalayan populations. **D)** Inferred cross-coalescent rate over time between the Sherpa and other non-Himalayan populations. **E)** Inferred cross-coalescent rate over time between the Lhasa and Himalayan populations. **F)** Inferred cross-coalescent rate over time between the Sherpa and Himalayan populations. A relative cross-coalescence rate of 0.5 is used as a heuristic approximation of the split time. Additional populations are plotted in Supplementary Figure 3.

To further understand how and when the observed population structure and patterns of ROHs appeared, we generated genome-wide genealogies^18^ to estimate coalescent to and reconstruct effective population sizes and separation histories. Our analysis reveals different population size trajectories in the Himalayan groups (Figure 2B). Some groups, notably the Lhokpu, Toto, Monpa, Layap, and to a lesser extent the Sherpa and Ghale, appear to have undergone a bottleneck in the past 10,000 years. The Lhokpu and Toto show the most significant decrease in population size (Ne = 1000-1500 in the past 3,000 years), consistent with the ROHs distribution. However, we note that recent inbreeding and population structure can result in bottleneck-like patterns in coalescent-based estimates^19^. Furthermore, some groups, such as the Magar, Bumthap, and Tshangla, have maintained stable population sizes over the past 10,000 years. The Lhasa population exhibits a notable increase in size starting around 5,000 ya, potentially indicating a period of population expansion or significant admixture. We then estimated relative cross-coalescence rates (rCCR) across populations and found that Lhasa separated from South Asian populations around 30,000 years ago (Figure 2C). However, when compared to certain East Asian populations (Tu, Yi, Naxi), we found that they begin to separate around 7,000 years ago (rCCR = 0.5), however these lineages show a gradual decrease in differentiation and do not fully separate (rCCR ∼ 0.25) over the past 5,000 years, indicating continuous gene flow until recent times. In contrast, when comparing the Sherpa to the same populations, a clearer split pattern appears (Figure 2D). The split time is approximately 10,000 years ago with the rCCR reaching almost zero in recent times, suggesting no or limited gene flow between these groups. When studying the Chetri, we find recent split time with the Pathan (3,000 years ago) and much older separation from the Sherpa (20,000 years ago), with low-levels of gene flow in the past 3,000 years (Supplementary Figure 3A).

Within the Himalayan populations, we also observe a contrast between the Lhasa and Sherpa. The Lhasa appear to have split from Kurtöp, Tshangla, and Bumthap relatively recently, within the past 2-5 thousand years, with ongoing gene flow until the present day (Figure 2E). The Sherpa, however, have an older split from these same populations, approximately 10,000 ya (Figure 2F). Other populations, such as the Layap, Ghale, Thakali, and Monpa, show a similar separation pattern to both the Lhasa and Sherpa, with a gradual split occurring around 10,000 ya. The Tharu also show similarly old split times (∼10,000 ya) from the Chetri and Magar, however recent gene flow is observed with both populations within the past 5,000 years (rCCR∼ 0.12, Supplementary Figure 3B).

### Migration and admixture in the Himalayas and surrounding regions

Next, we ran an ADMIXTURE^20^ analysis to estimate ancestry proportions within each individual in our dataset. We identified an ancestry component that is dominant in most Himalayan populations (Figure 3A, represented by the green colour). This component proportion is high not only in high-altitude populations like the Sherpa but also accounts for the entirety of the ancestry in mid- to low-altitude populations such as Monpa and Lhokpu. Integrating these results with our previous demographic analyses, the ADMIXTURE patterns likely reflect the effects of genetic drift exacerbated by bottlenecks, leading to pronounced differentiation. This is also evident, for example, in the CS Asian Kalash group, a known highly drifted population that appears to have a maximized ancestral component (Figure 3A), as previously reported^21^. Many Himalayan populations appear to have admixture from East and CS Asian populations. All Nepalese populations except Sherpa and Limbu have CS Asian ancestry (purple colour) which appears dominant in the Chetri population, reflecting their position on the PCA. Most Nepalese and Bhutanese populations have East Asian ancestry (blue colour) including the Tibetan Lhasa population which has a median of ∼2% East Asian ancestry, with one individual having ∼40% of East Asian ancestry reflecting ongoing admixture with lowland groups on an individual level (we subsequently remove this individual from all analyses involving the Lhasa population as a group). The ADMIXTURE plot further shows that lowland groups in East Asia, such as Tu, Yi and Naxi, as well as groups in Central Asia such as the Burusho, carry a proportion of Himalayan ancestry. Notably, this Himalayan component seems to be more pronounced in East Asia, with Yi and Naxi populations having approximately 50% of their ancestry derived from the Himalayan component, consistent with our coalescent-based analysis.

**Figure 3:**
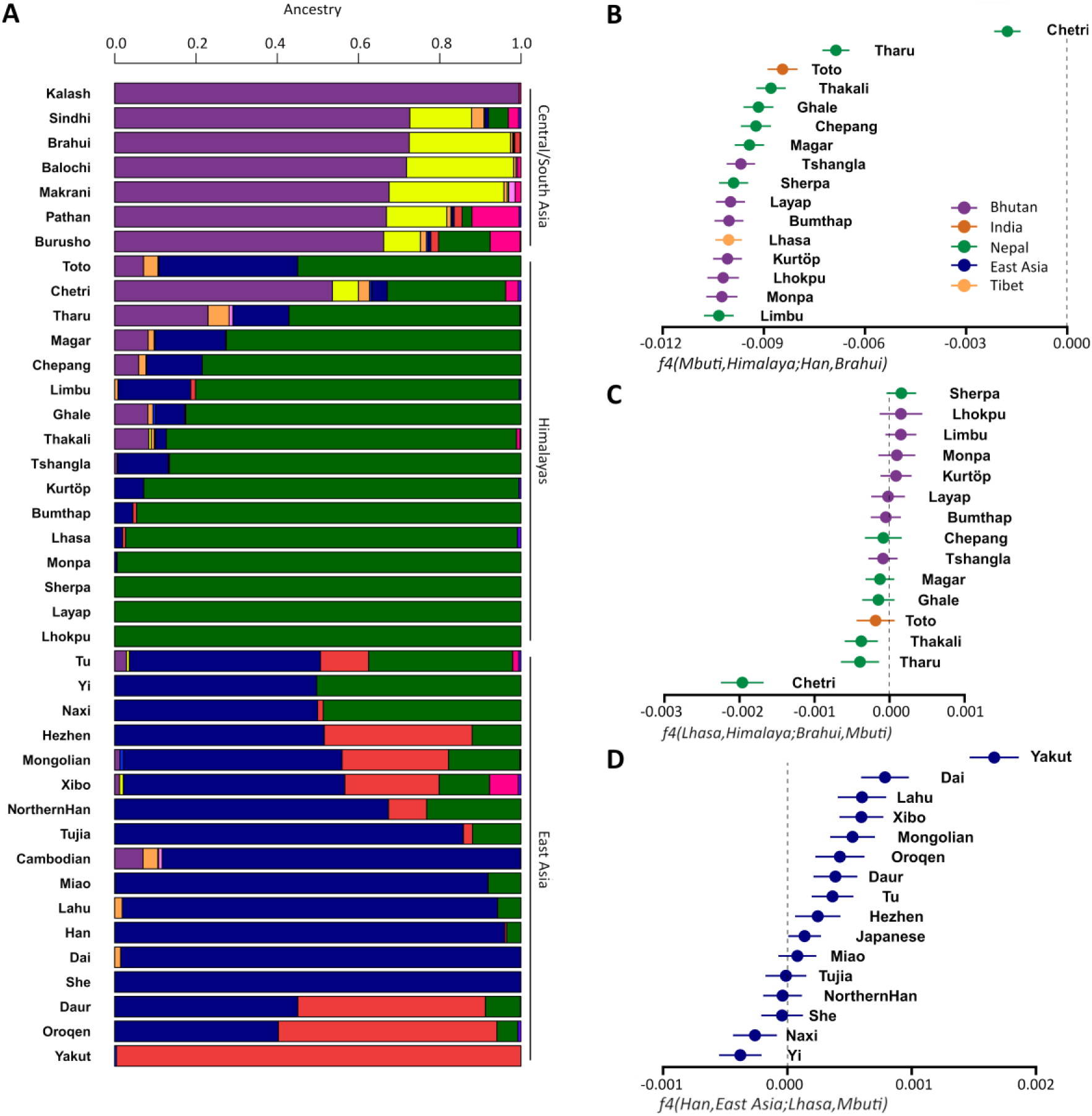
Migration and admixture in the Himalayas. **A)** ADMIXTURE analysis of Himalayan populations and lowland Central/Southern and East Asian populations. **B-D)** *f4*-statistics testing the relationships of the Himalayans to neighbouring populations. We plot the *f4* statistic value and ±3 standard errors.

Following our ADMIXTURE results, we examined population relationships using f4-statistics, testing *f4*(Outgroup, Himalayan; East Asian, CS Asian). This analysis resulted in significantly negative values (Z-scores < -3) for all tests suggesting that Himalayans are genetically closer to East Asians than to CS Asians (Table S2 and Figure 3B). Next, we computed *f4*(Outgroup, CS Asian; Himalayan, Tibetan) and found negative values when Himalayan was Chetri, Thakali, or Tharu (Z-scores <-3) or Ghale, Magar or Toto (Z-scores <-2) indicating increased genetic affinity of the Nepalese populations and Toto to CS Asians compared with Tibetans (Table S2 and Figure 3B). We then tested *f4*(Outgroup, Tibetan; East Asian, Han) and found significant negative values for East Asians Naxi and Yi reflecting the genetic affinity of these populations with Himalayans (Table S2 and Figure 3B), consistent with our earlier findings.

To understand the timeline of gene flow in the Himalayas, which led to the patterns described earlier, we employed a linkage disequilibrium (LD)-based statistic to estimate admixture dates^22,23^. We first tested for the arrival of CS Asian ancestry in the Nepalese populations Ghale, Magar, Chepang and Thakali. These populations share similar CS Asian ancestry proportions (approximately 6-9%) and cluster closely on the PCA plot. We therefore combined these populations into a single test group, which provided a larger sample size that enhances the accuracy of our estimates and reduces standard errors. Our estimates indicate that admixture in Nepal occurred approximately 51±7 generations ago (z=7.54, p=4.8e-14), or between 700 and 300 CE (Table S3).

Next, we tested the Nepalese populations Chetri and Tharu who have substantial South Asian ancestry (50% and 25% respectively - one Tharu individual had 68% SA ancestry). We did not obtain a significant z-score (>3) for the amplitude and the decay constant when we tested Tharu, probably because of the heterogeneity of admixture in the samples (CS Asian ancestry ranged from 22% to 68%). However, for the Chetri population we estimate admixture to have occurred 24±6 generations ago (z=4.03, p=5.7e-05) or between 1450 and 1100 CE (Table S3).

Similar to our tests for South Asian admixture in Nepal, we wanted to examine East Asian admixture in the Lhasa and Bhutanese populations. However, we did not obtain significant results, even when combining the Bhutanese Kurtöp, Tshangla and Bumthap populations or conducting individual tests. We propose that this outcome stems from background linkage disequilibrium and the genetic closeness between the reference East Asian populations and the test populations which potentially produces spurious weighted LD signal from recent shared demography.

We then investigated gene flow from the Himalayas towards the lowlands and therefore tested admixture in Burusho who have ∼13% Himalayan-related ancestry. When comparing the rCCR of the Burusho with Lhasa (Figure 2C), we find that they do not reach zero in the past 3,000 years, providing additional evidence of gene flow between these populations. We estimate admixture occurred 47±6 generations ago (z=8, p= 5.80E-16) or between 800 and 450 CE (Table S3).

We were interested in whether the admixture events we describe above might have been sex-biased, potentially involving the migration of different ratios of males and females between these regions. We tested this possibility by using qpAdm^24^ to estimate the admixture proportions, comparing the ancestry on the X chromosome with that on the autosomal genome. Because this test requires a set of outgroups that can differentiate between the sources of ancestry, we could apply it only to admixture events involving Himalayans and CS Asians but not Himalayans and East Asians. Our findings indicate that, for all the populations we tested, the proportions of ancestry on the X chromosome and autosomes were generally similar, except for the Chetri population, who appears to have experienced an excess of gene flow from CS Asian males compared to females (Table S4 and Figure 4A).

**Figure 4.**
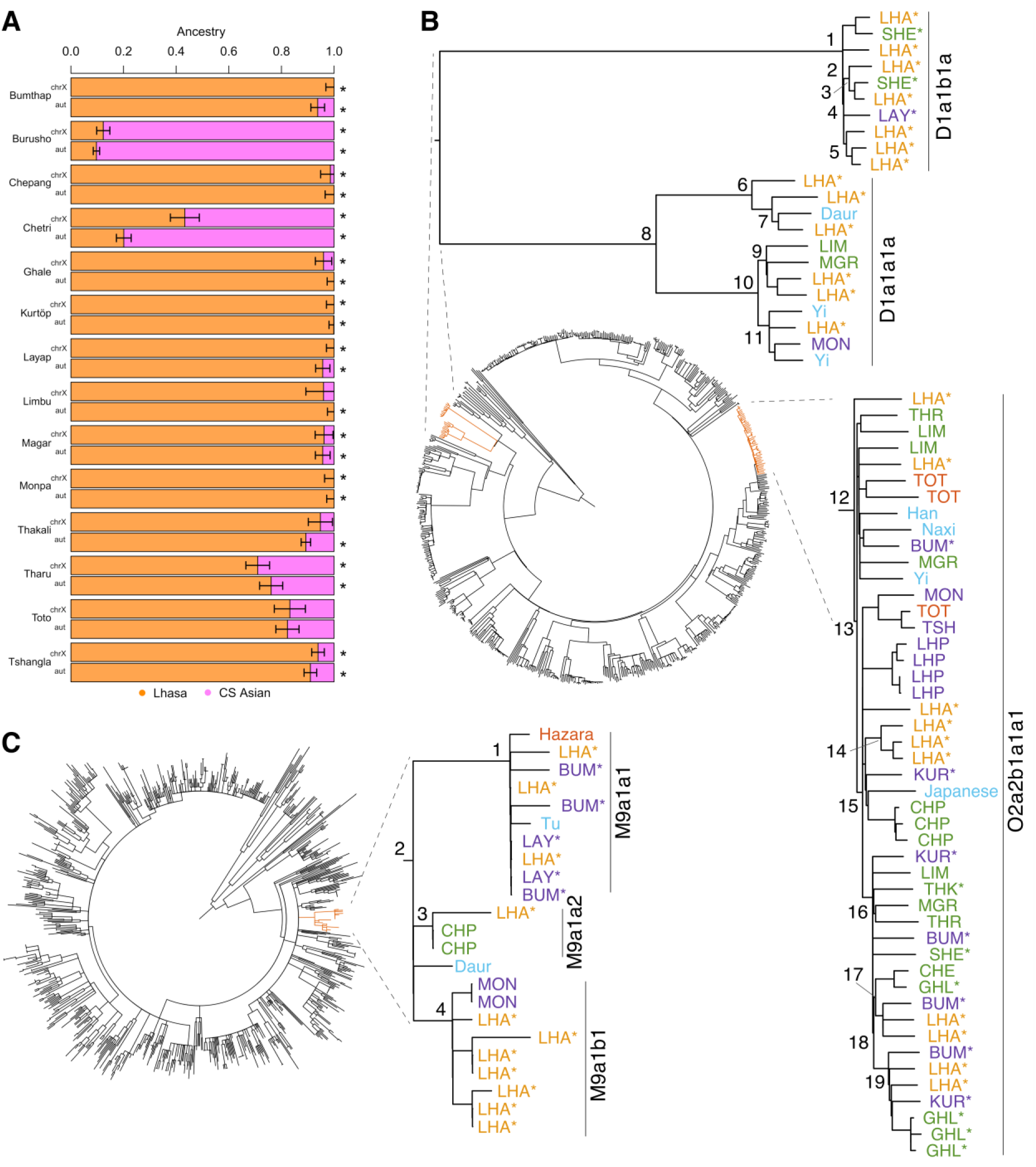
Sex biased migrations and uniparental lineages in the Himalayas. **A)** Estimates of admixture proportions in Himalayans comparing ancestry on the X chromosome with that in the autosomal genome, using Lhasa and Central/South Asian populations as sources of ancestry. A star indicates that the admixture model could not be rejected with a p-value threshold of 0.05. **B)** Worldwide Y- chromosomal maximum likelihood phylogenetic tree. The zoomed sections show the branches containing most of the Himalayan Y chromosomes. The numbers on the tree represent the nodes dated in Table S6. High-altitude individuals are indicated with a star (*). Population label colors indicate regions: blue for East Asia, light orange for Tibet, purple for Bhutan, and green for Nepal. **C)** Worldwide mtDNA maximum likelihood phylogenetic tree. The zoomed section shows the branches containing most of the Himalayan mtDNA haplogroups. The numbers on the tree represent the nodes dated in Table S6. High-altitude individuals are indicated with a star (*).

### Uniparental demographic history in the Himalayas

We constructed Y-chromosomal and mtDNA phylogenies using our dataset, revealing that both Y chromosome and mtDNA haplogroups in the Himalayas are a mixture of CS and East Asian lineages. The most common haplogroups for the Y chromosome are D1 and O2 with more than 82% of the males at high-altitude carrying the subclades D1a and O2a2b1a1a. The O2 lineage, common in South East Asian populations^25,26^, is the most widespread in the Himalayan region (Figure 4B, Table S5). Haplogroup D1 also forms two separate clades: D1ab1a1 and D1a1a1a. D1a1a1a and O2a2b1a1a show high-altitude specific clusters. By employing the ρ statistic, we estimated that the expansion time for the former occurred around 2,300-3,600 ya, while the latter expanded approximately 2,800-4,000 ya (Figure 4B, Table S6).

The most common mtDNA lineages in the Himalayas are M9 and D4 (Table S5). The D haplogroups are mostly found in Nepal and neighbouring lowland regions. The M lineage is widespread in South and East Asia^27^ with M4 and M5 found in lowland Himalayan populations. In contrast, high-altitude populations from Tibet, Bhutan and the Sherpa carry mainly the M9a lineage (Figure 4C), which has previously been associated with hypoxic adaptation in Tibetans^28^. We estimate the coalescence of the high-altitude M9a clusters to be around 3,400-5,600 ya.

### Archaic introgression and positive selection

*EPAS1* has been extensively investigated in Tibetans due to their possession of an introgressed Denisovan haplotype linked to high-altitude adaptation^13^. We sought to investigate whether similar selective pressures have manifested elsewhere in the Himalayas, particularly given our findings that Himalayan populations began diverging from each other within the last 10,000 years which is a time window that allows for the possibility of distinct populations undergoing differential selection over time. We used Patterson’s *D* and the related statistic 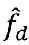 to test for introgression across Himalayan genomes^24,29^. We identified the introgressed *EPAS1* haplotype in all high-altitude Himalayan populations, with frequencies ranging from 40% (Bumthap, Kurtöp) to 80% (Lhasa) (Figure 5A). Notably, this haplotype is also present in lowland Himalayan populations and extends to Central Asian (Burusho, AF=11%) and East Asian (Yi, Tu, Naxi, AF=25%-38%) populations we earlier identified as admixed from Himalayans. This serves as additional evidence supporting gene flow from the Himalayas to the surrounding lowlands.

**Figure 5.**
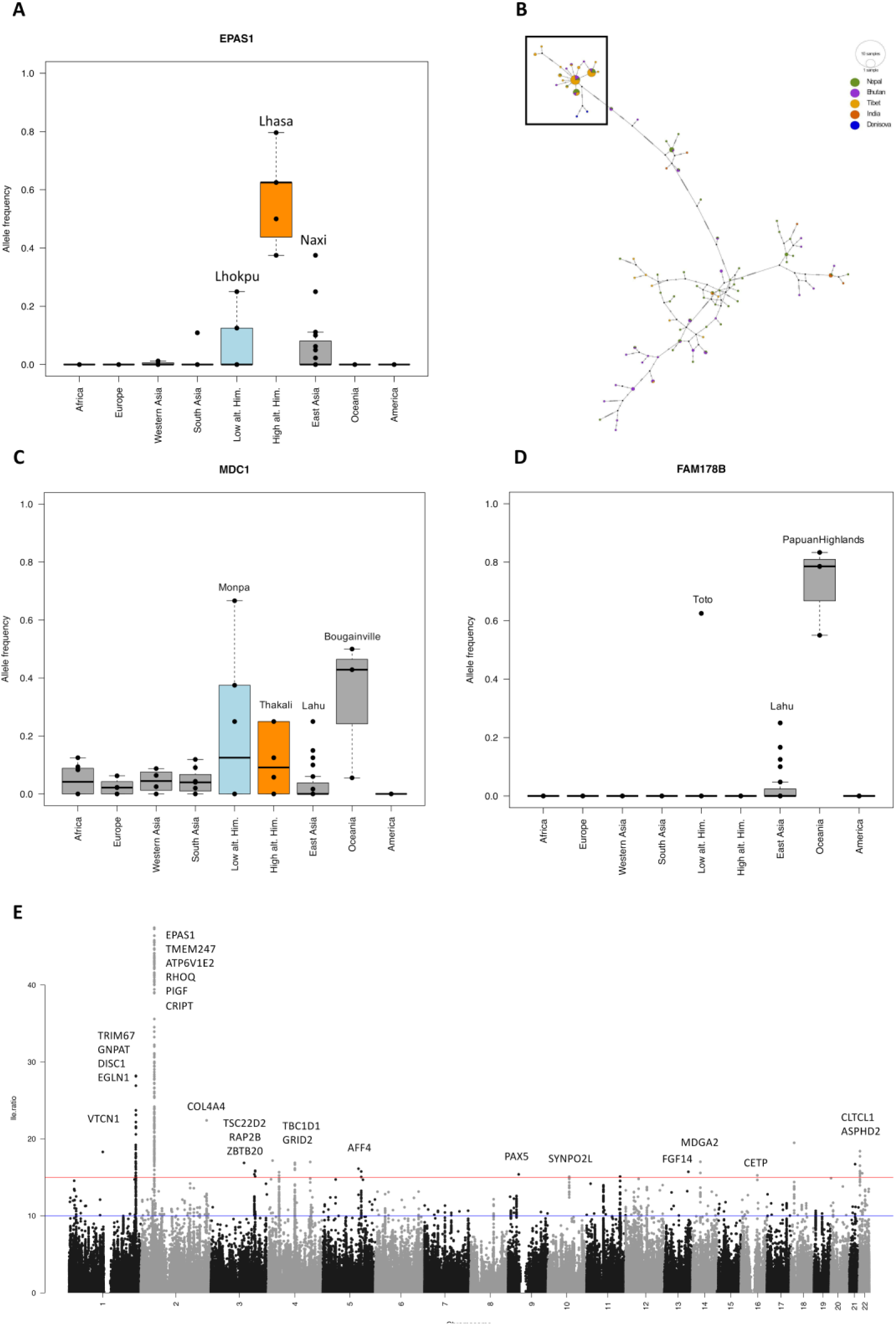
Archaic introgression and positive selection. **A)** The introgressed *EPAS1* haplotype of Denisovan origin appears at high frequency in all high-altitude Himalayan populations. **B)** Haplotype network of the introgressed *EPAS1* region in the Himalayan populations. The boxed area of the network contains the introgressed Denisovan haplotype. This cluster coalesces at 30,000-50,000 years ago. **C)** An introgression scan in the Monpa population reveals a haplotype at high frequency at *MDC1* in Monpa and Oceanians. **D)** An introgression scan in the Toto population reveals a haplotype at high frequency at *FAM178B* in Toto and Oceanians and at lower frequencies in South East Asian populations. **E)** Genome-wide selection scans show putatively adaptive variants in high-altitude Himalayans.

We ran a haplotype network analysis of the *EPAS1* haplotype region and found that high-altitude populations formed a star-like expansion together with the Denisovan haplotype. We then used the ƿ statistic to estimate that this cluster had a TMRCA of 30,000 to 50,000 ya (Figure 5B).

Beyond *EPAS1*, in the Monpa population who reside at a moderately-high elevation of 2,000 m, we identified a putatively Denisovan introgressed region on chromosome 6 spanning two adjacent genes, *NRM* and *MDC1* (Figure 5C). Interestingly, this haplotype is also present at high frequency in Oceanians (∼50%) and at lower frequency in other Himalayans and East Asians. Another potentially introgressed region was detected in the Toto population on chromosome 2 at *FAM178B* (AF=60%). This particular haplotype is also found at high frequency in Oceanians (AF=∼80%), moderate frequency in East Asians (AF=10-25%) but is absent in all other populations (Figure 5D).

Next, we conducted genome-wide scans for selection in the Himalayas, specifically seeking variants that are distinct in high-altitude populations compared to lowland groups, independent of archaic introgression. We used a method^30^ that accounts for population structure and admixture and identified variants that are putatively related to high-altitude adaptation in the Himalayas (Figure 5E, Table S7). The strongest signal was found at *EPAS1* and neighbouring genes such as *TMEM247, ATP6V1E2, RHOQ, PIGF, and CRIPT*, which have been previously implicated in high-altitude adaptation in Tibetans^31^. Another significant signal was observed at *GNPAT, TRIM67, and DISC1*, with *TRIM67* and *DISC1*, along with the nearby gene *EGLN1*, previously associated with hypoxia adaptation at high altitudes^32^. Additionally, we identified novel selection signals from variants in *TBC1D1, CLTCL1, PAX5, VTCN1, ZBTB20, SYNPO2L, CETP, and ASPHD2*. These genes harbour variants that are significantly differentiated between high-altitude populations and lowland groups.

## Discussion

The Himalayas are home to diverse ethnolinguistic human populations whose genetic relationships to each other and to global populations are not yet fully explored. The extreme elevations of the Himalayas present a unique and challenging environment for human populations and a geographic barrier that has influenced the movement of human populations throughout history, creating a unique and complex genetic diversity. To understand this genetic diversity, we analysed 87 new whole-genome sequences from 16 Himalayan populations and used them to give insight into population origin, structure, and genetic adaptations that characterize the human populations in this region.

We find that Himalayans diverged from East Asians around 10,000-12,000 ya, a period of climate warming and the advent of the Neolithic transition in several parts of the world, with many communities gradually shifting from a nomadic lifestyle to settled agriculture and animal domestication. By the mid-Holocene period, agriculture had become the primary livelihood for human populations in both the northern and southern regions of the Himalayas, though remnants of hunter-gatherer communities persisted in certain areas^33^. Our results indicate that high-altitude Himalayan populations began differentiating into present-day groups around this period, with some continued diverging into new groups until the last 2,000 years, mostly maintaining small population sizes throughout their history or experiencing bottlenecks.

We find surprisingly high levels of population structure within the Himalayas, with notably high *F*ST values, some of which are comparable to those observed between populations on different continents. This strong differentiation is likely due to high levels of genetic drift resulting from small effective population sizes over extended periods, coupled with limited gene flow among populations, which further increases genetic structure. Himalayans carry limited uniparental lineages, which can be an indication of a rapid expansion of a small founder population. The Himalayan subclades of Y-chromosome haplogroups O2 and D1 and the mtDNA M9 lineages all coalesce around 2,000-5,000 ya. This period coincides with the establishment of the first villages and the advent of agriculture, which facilitated permanent human occupation of the Tibetan Plateau after 3,600 ya^4^.

The Chinese Yi, Naxi, and Tu populations, located in close geographical proximity to the Tibetan Plateau with shared historical and cultural interactions, appear as the East Asian populations most closely related to the Himalayans in our dataset. The considerable gene flow detected from the Himalayas to these populations significantly contributes to this observed genetic affinity. Gene flow between East Asia and the Himalayas appears to have been bidirectional, with the East Asian gene flow being more prominent at mid to low altitudes. Populations residing above 4,000 m, such as the Sherpa and Layap, show no detectable impact from this East Asian gene flow. Interestingly, the Lhokpu, who live at low altitudes, share similar ancestry with these high-altitude groups and also do not appear to have received East Asian admixture. Nevertheless, our analysis reveals admixture in the Lhasa population, including an individual with half of their genome from lowland East Asian ancestry, most likely representing ongoing and recent interactions between the two regions.

Gene flow from the Himalayas to the lowlands extended westward, impacting CS Asian populations such as the Burusho, who reside in the nearby valleys located in the Karakoram Mountain Range in Pakistan. Our analysis indicates that the Burusho received Himalayan gene flow between 800 and 450 CE, a timeframe coinciding with the expansion of the Tibetan Empire into Central Asia. This period left a substantial impact on the cultural, political, and religious development of the region, and is supported by our genetic findings. Similar to the interaction with East Asia, gene flow between the Himalayas and CS Asia has been bidirectional. In our dataset, we observe that all Nepalese populations, except the Sherpa, have received gene flow from CS Asia, dating back to between 700 and 300 CE. This timeframe aligns with the rise and expansion of the northern Indian Gupta Empire^34^, which exerted political, cultural, and trade influence over major parts of Nepal, evidently leaving a genetic footprint as well.

While our findings indicate that Himalayans are genetically more closely related to East Asian populations than to CS Asians, there are exceptions, such as the Nepalese Chetri population. The Chetri population has substantial CS Asian ancestry, which we estimate to have been a result of admixture with the Nepalese between 1450 and 1100 CE. Historical accounts align with our genetic results, suggesting that Chetris migrated to Nepal from India during the 12th century following military invasions and instability in India^35^. Furthermore, our tests reveal that the Chetri migrations involved a significantly higher number of Indian males than females. On the other hand, for the Tharu population, which also has a strong genetic affinity to CS Asians, our findings indicate a similar contribution from Indian women and men to the ancestry of the Tharu population, aligning with historical accounts of the Tharu descending from Rajput families escaping military invasions^35^.

Previous studies showed that Tibetans and other Himalayans carry an introgressed Denisovan haplotype associated with high-altitude adaptation^13,36^. Our results confirm the widespread distribution of the introgressed *EPAS1* region among high-altitude Himalayan populations, highlighting its critical role in survival within this challenging environment. Our estimated time to the most recent common ancestor for the Denisovan *EPAS1* cluster is approximately 30,000-50,000 ya, suggesting it could have been carried by hunter-gatherer groups inhabiting the region before disseminating with populations that later expanded to high altitudes. It has been proposed that this haplotype remained selectively neutral for a significant period following the introgression event, with selection pressures only beginning around 9,000 ya^37^.

We did not find evidence of further selection on Denisovan introgressed regions in high-altitude Himalayans in general. However, we detected a Denisovan haplotype with a high frequency in the Monpa population, spanning *NRM* and *MDC1*, genes associated with genome stability and lymphocyte counts. This haplotype is also prevalent in Oceanian populations, who carry extensive Denisovan introgression. We note that this genomic region lies outside the 1000 Genomes Project (1KGP) strict mask for short-read sequencing accessibility, emphasizing the need for confirming the results in the future through long-read sequencing.

We identify a third potentially introgressed region on chromosome 2 at *FAM178B*, exhibiting a high frequency within the Toto population and also prevalent among Oceanians.

Intriguingly, this region was identified previously as of Denisovan origin and under selection in the breath-hold diving Bajau people of Indonesia^38^. *FAM178B* appears important for adaptation to hypoxia, playing a role in blood pH regulation and preventing carbon dioxide accumulation. However, the link to the Toto population is enigmatic as today they inhabit lowlands in India. The preferentially endogamous Toto population represents a remnant of a larger number of Toto settlements formerly scattered throughout the Western Duars as far west as the Teesta, that were assimilated by more populous Bodo-Koch groups. Other than the village of Totopara, former Toto territory has been swallowed up by tea estates. The Toto language is part of the Dhimalish subgroup, which shows distant affinity with Bodo-Koch languages^39^. Here, we present two plausible scenarios to interpret the results: 1- The Toto population might have originated from or regularly visited nearby high-altitude regions in Bhutan, where an introgressed region at *FAM178B* from an ancestral population enhanced their survival. This model requires the loss of this introgressed region from all other tested neighbouring populations. 2- Alternatively, the Toto population could have experienced significant gene flow from Southeast Asia, where the frequency of introgressed variants is approximately 25% in our dataset in populations like Lahu and Cambodians. Subsequently, these variants might have increased to high frequency in the Toto population due to drift or adaptation as they migrated to settle near the mountains of Bhutan. Notably, the Toto exhibit the highest genome-wide proportion of East Asian ancestry among all Himalayans in our dataset and cluster closely with Cambodians on the PCA.

Our genome-wide scans for selection, conducted without utilizing archaic genomes, also identify variants at *EPAS1* as having the strongest signal. The genomic region extends to adjacent genes, including *TMEM247*, *ATP6V1E2*, *RHOQ*, *PIGF* and *CRIPT*, all of which have been previously recognized as candidates for adaptation in Tibetans through allowing biological adjustments to changing oxygen levels^31^. Another strong signal appears on chromosome 1 in a region spanning *GNPAT*, *TRIM67* and *DISC1*; the latter two genes were previously found to be associated with high altitude adaptation to hypoxia, along with the neighbouring gene *EGLN1*^32^. *GNAPT* was previously associated with skin pigmentation in Tibetans for UV protection^40^. We identify a novel selection signal on variants located on chromosome 4 in *TBC1D1*. In mice, *TBC1D1* plays a crucial role in glucose and lipid utilization, as well as influencing energy substrate preference in skeletal muscle^41^. Notably, individuals residing at high altitudes demonstrate lower fasting glycemia and enhanced glucose tolerance compared to those in lowland areas, coupled with emerging evidence pointing to a reduced prevalence of obesity and diabetes at higher altitudes^42^. Furthermore, this gene has been associated with cold adaptation in chickens, likely through its role in regulating energy homeostasis^43^. We also find a strong selection signal on chromosome 22 at *CLTCL1*, a gene responsible for directing the production of the CHC22 protein which plays an important role in regulating a glucose transporter in both fat and muscle cells^44^. We find additional putative adaptation signals in: *PAX5* associated with B-cell development^45^, *VTCN1* and *ZBTB20* implicated in the regulation of T cells^46,47^, *SYNPO2L* which plays a role in the structural development and function of cardiac myocytes^48^, *CETP* responsible for mediating the transfer and exchange of cholesteryl ester and triglyceride in addition to an important role in HDL metabolism^49^, and *ASPHD2* which enables dioxygenase activity and metal ion binding activity^50^. These genes collectively harbour variants that exhibit significant differentiation in high-altitude populations compared to lowland Himalayans and neighbouring populations.

These genetic variations could have played a role in shaping aspects of cellular and metabolic functions, potentially contributing to enhanced survival in the challenging conditions of the Himalayas.

In conclusion, our research significantly advances our comprehension of genetic diversity, origins and adaptations within human populations residing in the Himalayas. While prior studies concentrated on the genetics of a few ethnic groups in Tibet, our investigation introduces insights into little-explored populations from the region and on a whole-genome level. This broader perspective enhances our understanding of the complex genetic diversity and adaptive processes shaped by the distinctive environmental conditions of the Himalayas.

## Material/Subjects and Methods

### Samples and dataset

We sequenced 87 individuals from 16 Himalayan populations using the Illumina HiSeq X Ten platform at the Wellcome Sanger Institute to an average depth of ∼34x. The dataset included eight populations from Nepal, six from Bhutan, one from North India, and Tibetans from China. Seven of the Himalayan populations reside at altitudes of 2,500 meters or above (Table S1). For some of our analyses, we categorize these as ‘high-altitude’ populations. These samples, except the Tibetans, were collected as part of the ‘Language and Genes in the Greater Himalayan Region’ project^51^. Tibetans were collected previously to study human demography in East Asia^52^.

### Variant calling and filtering

We mapped raw sequence reads to the human reference assembly GRCh38 (GRCh38_full_analysis_set_plus_decoy_hla.fa) using BWA mem^53^. Base quality score recalibration, indel realignment and duplicate removal were performed using the Genome Analysis Toolkit (GATK)^54^ v4.1. One genomic VCF (gVCF) per sample was generated using GATK HaplotypeCaller and then joint-called with Human Genome Diversity Project (HGDP)^15^ single sample gVCFs using the GATK GenotypeGVCFs tool. A set of standard hard filtering parameters for autosomal SNPs and INDELs was applied to the joint call set^15^: 1) for each sample, any genotype with a GQ or RGQ value equal or lower than 20 or a depth of coverage (“DP” annotation) equal to or greater than 1.65 times the genome-wide average coverage for the sample was set to missing; 2) the bcftools^55^ v1.7 “ExcHet” filter tag was used to filter out variant sites with an ExcHet value equal to or larger than 60 (equivalent to an excess heterozygosity p-value of 10^-6^); 3) we restricted our analyses to variant sites in the masked region of the genome from the 1000 Genomes Project^56^ accessibility strict mask for GRCh38 (version 20160622). From this mask, we also removed any regions of the primary GRCh38 assembly which have alternative loci or patch scaffolds, as defined by the NCBI assembly resource for the GCA_000001405.15_GRCh38 entry. Variants with missingness greater than 0.2 were filtered out. This resulted in a final dataset of autosomal ∼70M SNPs and ∼9M indels. Only SNPs were used for demographic analyses.

Variant calling and filtering for the Y chromosome and mtDNA was performed using a different approach. The call set for the Y chromosome was restricted to the commonly used ∼10.3 Mb region accessible by short-read sequencing^57^. All Himalayan samples were confirmed to be males by analysing the average read depth across the accessible regions on both X and Y chromosomes using GATK v4.1 DepthOfCoverage and found to be similar. All-sites joint calling was performed using BAM files for all male samples of HGDP and Himalayans using bcftools v1.8 mpileup and call flags for the ∼10.3 Mb callable regions, setting the ploidy argument to 1, using –m (multiallelic-caller) and setting the minimum base quality (-Q) and the minimum mapping quality (-q) to 20. The raw calls were filtered for SnpGap 5 (removal of single nucleotide variants within 5 bp of an indel), followed by removal of indels. Genotypes from samples with an average read depth of over 12 on the Y chromosome were filtered with a minimum read depth of 3. For samples with lower mean read depth, the minimum was 2. If multiple alleles were supported by reads, then the chosen allele needed support from at least 85% of the reads; otherwise, the genotype was marked as missing. Sites with 3% or more missing calls across samples were excluded. After filtering a total of 709 samples and 10,292,707 sites remained, including 56,002 variant sites. The haplogroups for Y chromosomes were defined using yHaplo^58^. The identified terminal marker for each sample was used to update the haplogroup name to correspond to the International Society of Genetic Genealogy (ISOGG, https://isogg.org) nomenclature v14.255.

All-sites joint calling of mtDNA was performed similarly to the Y chromosome using bcftools mpileup. Mitochondrial sites showing evidence of heteroplasmia were removed for downstream analyses. For the construction of mtDNA phylogeny and dating only the coding region was used. The haplogroups for mtDNA were defined using HaploGrep v2.0^59^.

### Population structure analyses

We computed a PCA using smartpca v16000 from the EIGENSOFT package^60^ with parameters numoutlieriter: 0. The analysis included Himalayans, CS Asians, and East Asians. We retained SNPs with a Minor Allele Frequency (MAF) greater than 0.05 and carried LD pruning with PLINK^61^ using --indep-pairwise 200 25 0.4. Figures were plotted using R^62^. Map in Figure 1B was plotted using ggmap^63^. We also calculated pairwise *F*ST using smartpca v16000 on a set of 1.3M SNPs that were ascertained as polymorphic in archaic genomes as they produce more accurate estimates^15^.

We ran unsupervised ADMIXTURE^20^ analyses from K=6 to K=20 using the complete dataset, which includes many populations from other global regions. Similar SNP filtering criteria to those employed in the PCA were applied. We show the results for K=11 which had the lowest cross-validation error.

We used qpDstat v980 from the ADMIXTOOLS package^24^ with parameter f4mode: YES to test genetic affinity between Himalayans, and CS and East Asians. From the same package we used qpAdm v1520 to test for sex-biased migrations by contrasting ancestry proportions on the X chromosome and chromosome 13 (representing the autosomal genome and with comparable size to the X). We used the following outgroups: Mbuti, Japanese, Basque, Papuan Sepik, Pima, Surui, Yakut, Bedouin and Kalash.

We estimated admixture dates with MALDER v1.0^22,23^ using parameters mindis: 0.005 and binsize: 0.001 and reference populations Lhasa, Sherpa, Kalash, Brahui, Pathan and Balochi to test admixture in Nepal and CS Asia. We used reference populations Lhasa, Sherpa, Han, Lahu, Dai and She to test admixture in Bhutan and East Asia. We set genetic positions using the HapMap genetic map supplied with Eagle v2.4.1^64^ (genetic_map_hg38_withX.txt.gz). We show the date from population pairs that have the lowest p-value and assume one generation time is 29 years^65^.

### Coalescent analysis

Since the accuracy of coalescent-based inference depends on phasing quality, we evaluated different phasing strategies for the HGDP+Himalaya joint-called dataset by comparing the results to an experimentally-phased Han sample published in the HGDP WGS dataset^15^. We tested the following approaches: 1-Reference-free phasing using Eagle v2.4.1. 2- Reference-based phasing using the 1KGP high-coverage panel^66^ with Eagle, followed by Beagle v4.1^67^ with the option usephase=true to phase the remaining variants not present in the reference panel, using information within the dataset and the previously generated phased data as a scaffold. 3- Similar to the second approach, but utilizing the phased 1KGP+HGDP reference panel^68^.

By comparing the experimentally-phased VCF of HGDP00819 with the statistically-phased sample within our joint-called dataset on a set of heterozygous variants extracted from chromosome 8, we found the third phasing strategy described above to yield the lowest switch error rate (0.08%, excluding singletons as they cannot be statistically phased). Subsequently, we proceeded with this phased dataset to create genome-wide genealogies using RELATE v1.2.1^18^, employing 164 haplotypes (6 per population) and excluding any outliers identified in the PCA and ADMIXTURE analyses. We used a generation time of 29 years^65^, a mutation rate of 1.25×10^-8^, and diploid effective population size of 30,000^18^.

### Runs of homozygosity (ROHs)

We pruned the joint-called dataset using PLINK v1.90b6.24 with the options --geno 0.05 -- indep-pairwise 50 5 0.4 --maf 0.05 and subsequently ran --homozyg which identified ROHs segments 1Mb and larger.

### Phylogenetic reconstruction and dating for Y chromosome and mtDNA

Maximum likelihood phylogenies were constructed using RAxML v8.2.12^69^ with the GTRGAMMA substitution model. The FigTree software (http://tree.bio.ed.ac.uk/software/figtree/) was used to visualise the tree with midpoint rooting.

The ages of nodes of interest in both the Y and mitochrondrial phylogenies were estimated using the ρ statistic^70^ as described previously ^71^. In short, the all-sites vcf files were used to calculate pairwise divergence estimates. If multiple samples were present in a given clade, then the per-pair divergence estimates were averaged across them. The divergence estimates (mutations per site) were converted to units of years using a point mutation rate of 0.76×10^−9^ (95% CI: 0.67-0.86×10^-9^) mutations per site per year^72^ for the Y chromosome and 1.57×10^-8^ (1.17-1.98×10^-8^ 95% highest posterior density (HPD)) mutations per site per year for the mitochondrial coding region^73^. The 95% confidence intervals of the divergence times were estimated using the 95% confidence and HPD intervals of the mutation rates.

### Archaic introgression and positive selection

We searched for Denisovan introgressed regions on the genomes of the Himalayans using Dsuite^74^ with parameter -w 20,10 representing the window of tested SNPs and the moving step. We tested *D*(Han, Himalayan; Denisovan, Chimp) and filtered out results that did not have both Patterson’s *D* and 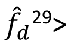 0.8. We then annotated the remaining positions using the Variant Effect Predictor (VEP) v107 and calculated allele frequencies for each position using bcftools v1.10.2. When discussing allele frequency of a haplotype in the text and in Figure 5, we refer to the lowest AF found at any position in the haplotype.

We used the Ohana software^30^ to identify signals of adaptation in high-altitude Himalayans. We began with an inference of global ancestry and covariance structure of allele frequencies among CS and East Asians, and high-altitude Himalayan populations, specifying ancestry components K=3. Iterations continued until the likelihood improvement fell below 0.001 (-e 0.001). Then, we chose markers with a log-likelihood ratio (LLRT) of ≥ 10, prioritizing those with the highest allele frequency among high-altitude Himalayan populations. The results were visualized through a Manhattan plot generated using qqman^75^.

### *EPAS1* network analysis

We generated the *EPAS1* network with POPART^76^. Using the median-joining network, the root of the human introgressed haplotype was assigned as the node where the Denisovan haplotype branch met the human introgressed haplotype branch. The mean number of mutations between this node and each human haplotype was counted (ρ statistic) and converted into a time in years assuming a mutation rate of 0.5 x 10^-9^ mutations per bp per year^77^.

## Data Availability

Aligned BAM files are available from European Genome-phenome Archive under accession number EGASxxxxxxx.

## Supplementary Material

Three figures and seven tables.

## Acknowledgments

We thank the sample donors who made this research possible. We thank the Wellcome Sanger Institute sequencing facility for generating data. Computations were performed using the University of Birmingham’s BlueBEAR HPC service.

---

**Figure S1.**
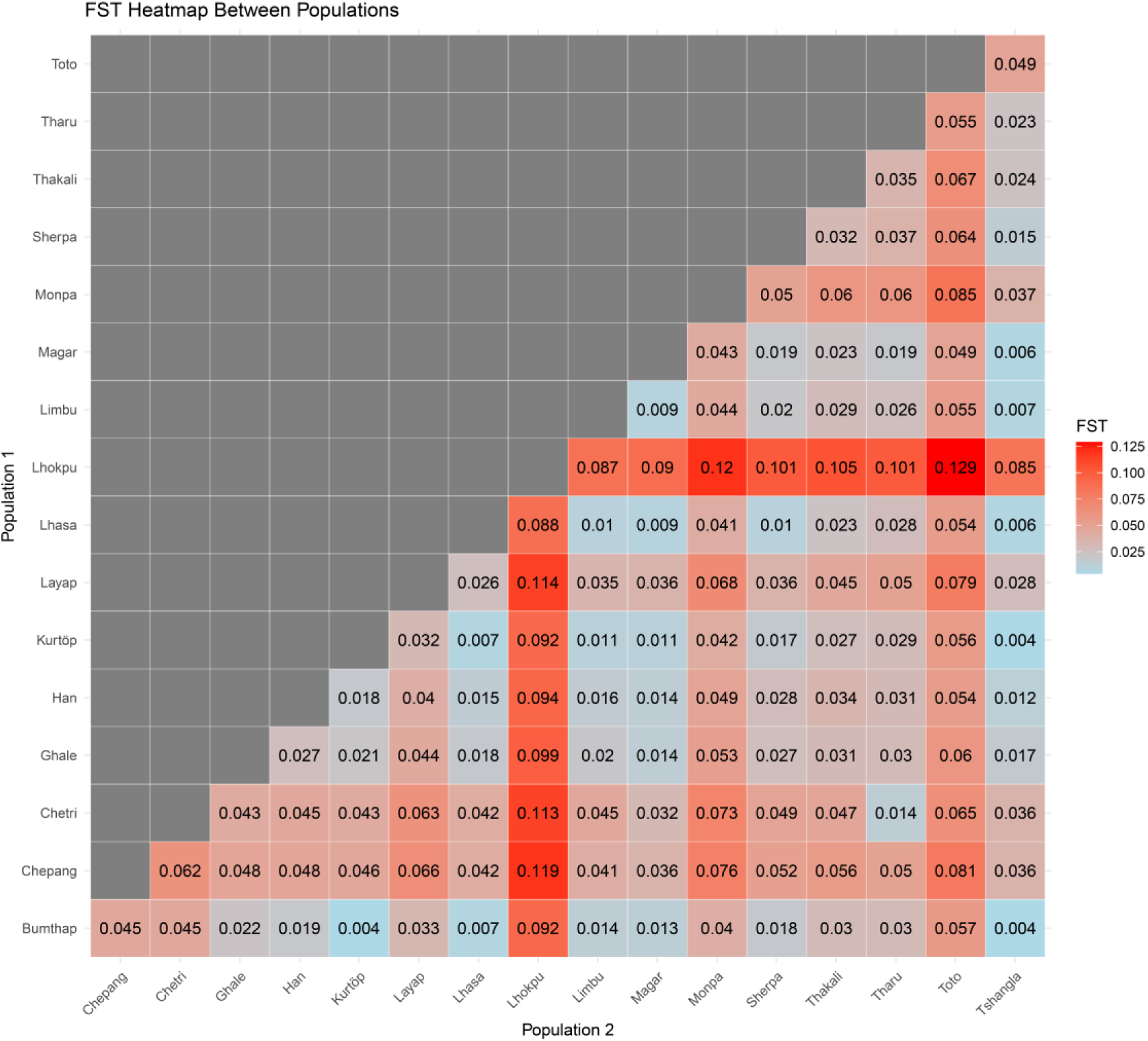
Pairwise *F*ST Values for Himalayan populations investigated in this study in addition to the Chinese Han.

**Figure S2.**
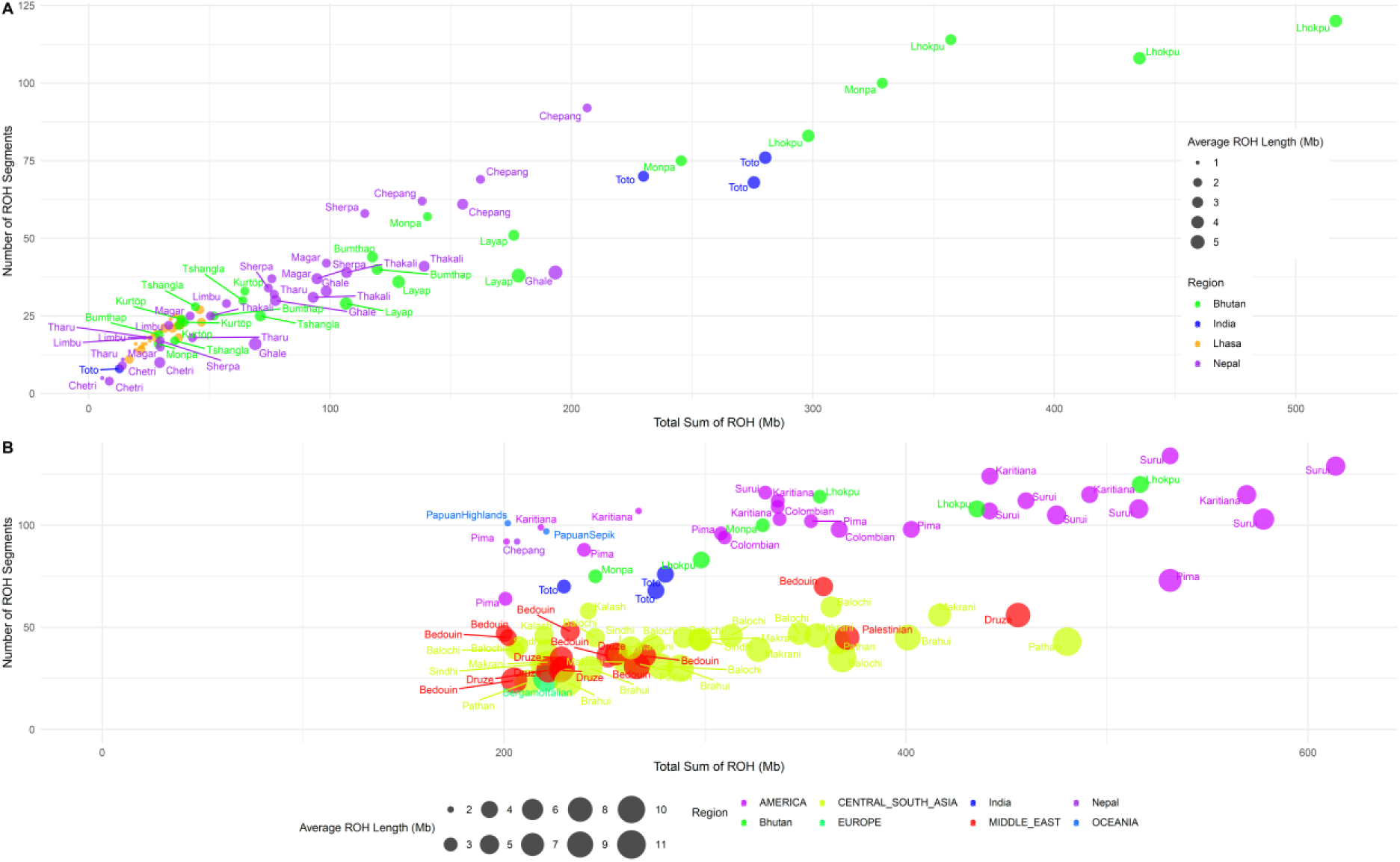
Runs of Homozygosity distribution in Himalayan and global populations. **A**) ROHs distribution in the Himalayas for all samples sequenced in this study. **B)** Samples with sROH > 200 Mb from the HGDP and Himalayan samples. Note that Middle Eastern and Central and South Asian populations have larger Average ROH length.

**Figure S3.**
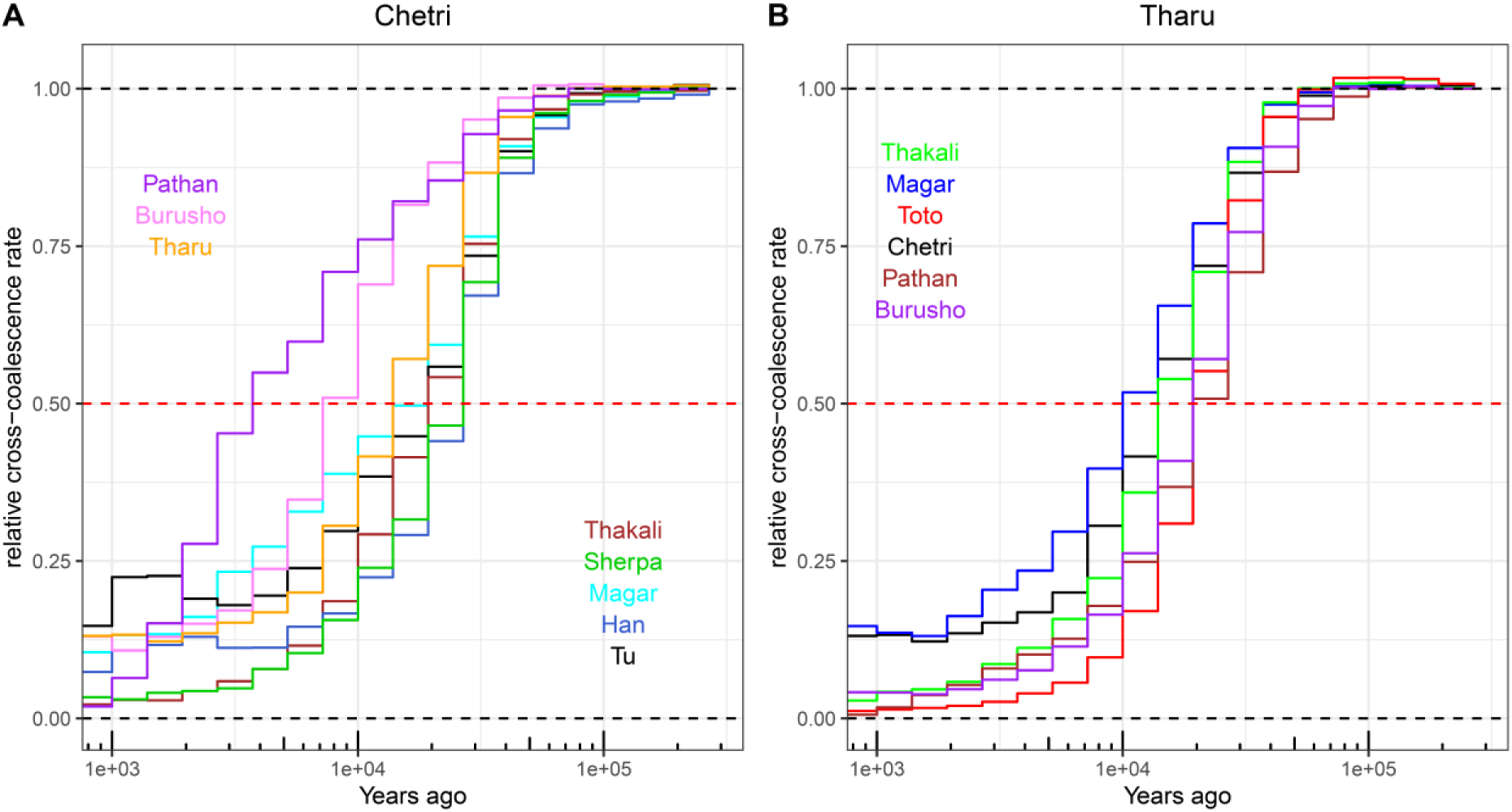
**Coalescent-based separation time history analysis**. **A**) Chetri and **B**) Tharu.

